# scATAC-pro: a comprehensive workbench for single-cell chromatin accessibility sequencing data

**DOI:** 10.1101/824326

**Authors:** Wenbao Yu, Yasin Uzun, Qin Zhu, Changya Chen, Kai Tan

## Abstract

Single cell chromatin accessibility sequencing (scCAS) has become a powerful technology for understanding epigenetic heterogeneity of complex tissues. The development of several experimental protocols has led to a rapid accumulation of scCAS data. In contrast, there is a lack of open-source software tools for comprehensive processing, analysis and visualization of scCAS data generated using all existing experimental protocols. Here we present scATAC-pro for quality assessment, analysis and visualization of scCAS data. scATAC-pro provides flexible choice of methods for different data processing and analytical tasks, with carefully curated default parameters. A range of quality control metrics are computed for several key steps of the experimental protocol. scATAC-pro generates summary reports for both quality assessment and downstream analysis. It also provides additional utility functions for generating input files for various types of downstream analyses and data visualization. With the rapid accumulation of scCAS data, scATAC-pro will facilitate studies of epigenomic heterogeneity in healthy and diseased tissues.

## Background

Chromatin accessibility is a strong indicator of the activities of functional DNA sequences. Recently, multiple experimental protocols have been developed to profile genome-wide chromatin accessibility in single cells, including the Assay of Transposase Accessible Chromatin with high throughput sequencing (scATAC-seq) [1], Single-cell Combinatorial Indexing ATAC-Seq (sci-ATAC-seq) [2], single-cell transposome hypersensitivity site sequencing (scTHS-seq) [3], and droplet-based single-cell combinatorial indexing ATAC-Seq (dsciATAC-seq) [4]. In this paper, we collectively define data generated with different experiment protocols as single cell chromatin accessibility sequencing data, or scCAS data. Application of these assays have helped to understand the epigenetic heterogeneity across cell populations in complex tissues during normal development and pathogenesis, including adult mouse tissues [5], forebrain development [6], hematopoietic differentiation and leukemia evolution [7,8], T cell development and exhaustion [9].

In contrast to the rapid growth of scCAS data, bioinformatic tools for scCAS data analysis are critically lacking. The majority of existing analytical tools lack comprehensiveness in their ability to process scCAS data. Both *chromVar* [10] and single-cell regulome analysis toolbox, *SCRAT* [11] work with preprocessed data and only report loss or gain of chromatin accessibility on a set of predefined genomic regions, which ignores a large amount of information in the data. Detection of cell-type specific difference in chromatin accessibility, *Detin* [12], single cell accessibility based clustering, *scABC* [13] and *cisTopic* [14], focus on identifying cell populations and/or differential accessible regions given the processed data like bam files or in peak-by-cell count matrix.

Single-cell ATAC-seq analysis tool *Scasat* [15] and *scitools* [16] are the only published software for comprehensive analysis of scCAS data. However, Scasat is developed in the Jupyter notebook environment. Although it is interactive, the programming codes are hard to standardize and reuse and users need to customize the analysis step by step. Furthermore, *Scasat* binarizes raw peak-by-cell count matrix, which ignores the differences among accessible regions and thus may lead to loss of valuable information for downstream analysis. In addition, *Scasat* does not provide summary reports for either data quality assessment or downstream analysis. *scitools* only works for sciATAC-seq and is not well documented. Another tool *SnapATAC* [17] also binarizes the raw count data and cluster cells based on bin-by-cell count matrix. *Cellranger-atac* by 10x Genomics (https://www.10xgenomics.com) is another comprehensive tool but only works with data generated using the 10x platform, and the software code is not open source. Additionally, some key analytical modules of *Cellranger-atac* are not flexible and do not use state-of-the-art algorithms. For example, the peak calling task does not use state-of-the-art algorithms, such as Model-based analysis for ChIP-Seq 2, *MACS2,* [18], resulting in many problematic peaks.

Here, we present a comprehensive and open-source software package for quality assessment and analysis of single-cell chromatin accessibility data, scATAC-pro. It provides flexible options for most of the analytical modules with carefully curated default settings. Summary reports for both quality assessment and downstream analysis are automatically generated. Interface to an interactive single-cell data visualization tool *VisCello* [19] is also provided.

## Results

### scATAC-pro provides flexible choices of methods for many analytical tasks

The overall workflow of scATAC-pro is depicted in Figure 1. We provide at least two alternative methods for all data processing and analytical modules. There are several reasons to have alternative methods for scCAS data processing and analysis. First, for certain tasks, many tools exist that perform equally well but have different levels of trade-off between mapping accuracy and mapping speed [20]. For example, for read mapping, several popular aligners exist, including BWA [21], Bowtie [22] and Bowtie2 [23]. Users can choose among those aligners based on different goals. For cell calling, cells can be called by filtering low-quality barcodes [1,2,5,8] or by using model-based approaches (e.g. *cellranger-atac*). Each class of methods has its pros and cons that can be tailored towards the goals of the analysis.

**Figure 1.**
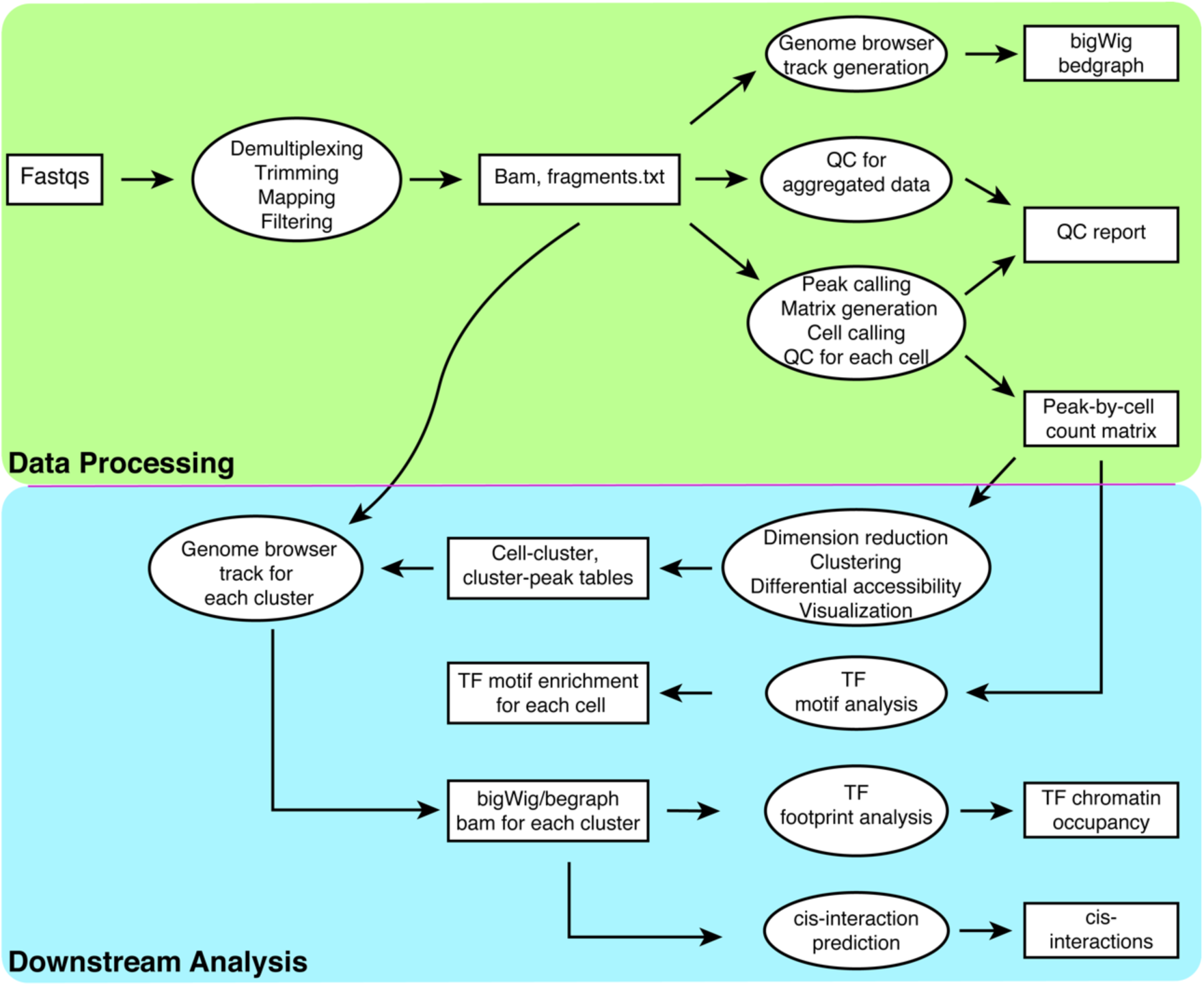
The scATAC-pro workflow. The workbench consists of two units, data processing unit and downstream analysis unit. Modules of the data processing unit include demultiplexing and adaptor trimming of the raw reads, followed by mapping of reads to the reference genome, filtering of low-quality reads. Aggregated reads are used for generating genome browser tracks, peak calling and cell calling. Quality check (QC) reports are generated based on both aggregated data and single cell data. Modules of the downstream analysis unit consists of dimension reduction, clustering, differential accessibility between different cell populations, genome browser track generation, TF binding motif and footprinting analyses, and prediction of chromatin interactions. Most modules provide more than one analytical methods.

Second, in the cases where a general-purpose and top-performing method exists, such as *MACS2* [18] for peak calling, there are other methods that are more suitable for specific tasks. For example, the genome wide event finding and motif discovery algorithm, *GEM* [24], was shown to have better performance in identifying peaks overlapping with transcription factor binding sites (TFBS) [25]. In this case, the users might prefer *GEM* over *MACS2*.

Third, often times there is a need for methods that can address dataset-specific characteristics. For example, it is more appropriate to binarize the peak-by-cell count matrix if the sequencing depth is shallow. Ideally, binarization of the count matrix should be avoided because non-binarized counts provides differential accessibility information. Therefore, we provide methods that work on binarized and non-binarized peak-by-cell count matrices (See details in Methods). Another example is clustering, a critical task for understanding heterogeneity in a cell population. To help selecting better clustering methods, we conducted a benchmarking study using simulated data. The compared methods include *scABC, chromVAR, cisTopic*, Latent Semantic Indexing or *LSI* [2], *SCRAT*, and Louvain algorithm implemented in *Seurat* v3 [26]. Based on the benchmarking result (Supplementary Figure 2), we choose *cisTopic* and *Seurat* as the methods for the clustering module. Note that *cisTopic* binarizes the peak-by-cell count matrix while *Seurat* does not.

**Figure 2.**
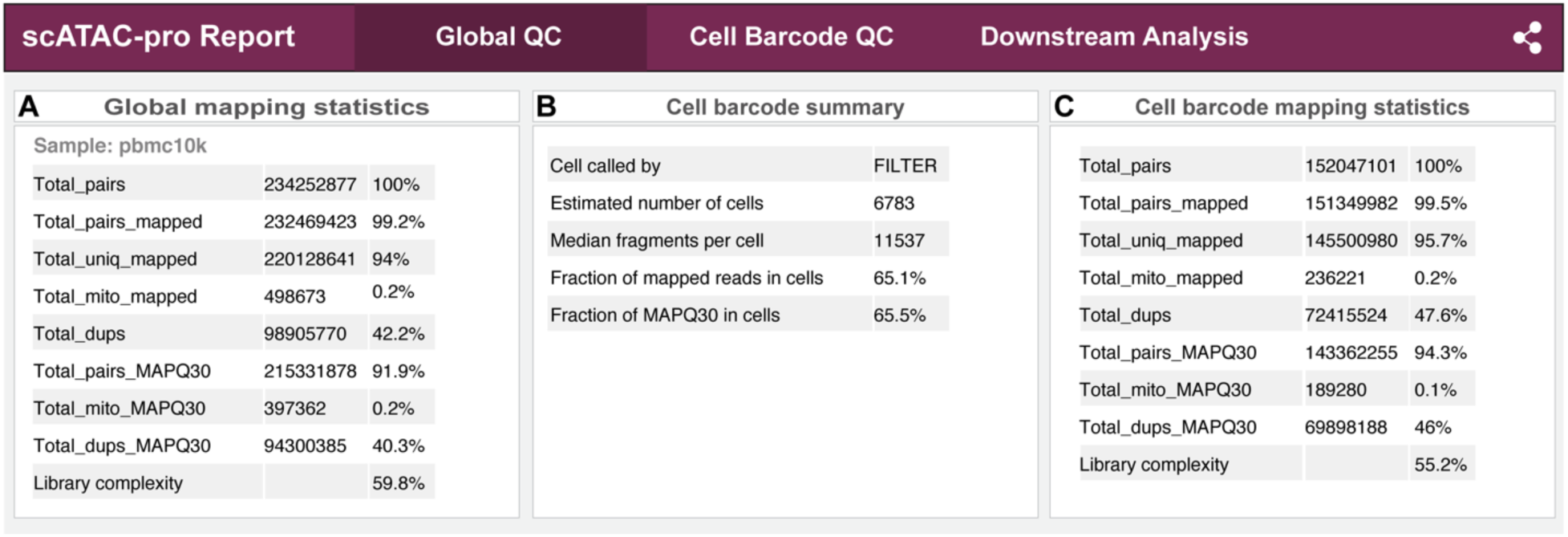
Summary statistics for read mapping, library complexity, and cell calling. scATAC-Seq data of human peripheral blood mononuclear cells (PBMCs) was used for illustration purpose. Global mapping statistics are based on all data (**A**). Cell barcode mapping statistics are based on called cells (**B, C**). MAPQ, mapping quality score.

### scATAC-pro provides carefully evaluated default settings for all modules

Method and parameter choice makes a big difference in the result of several analytical modules. We therefore provide a set of carefully evaluated default settings for each analysis module. We discuss the setting for these modules as follows (see Methods for details).

#### Read mapping

Because of its balance between mapping speed and accuracy [27], especially for paired-end sequencing data, we choose BWA (specifically bwa-mem) as the default read aligner.

#### Peak calling

Peak calling is usually done on aggregated data across all barcodes. Such an approach fails to identify peaks that only appear in rare cell populations. We implemented a two-step strategy, similar to the idea used by [5]. We first segment the genome into 5 kb bins and generate a bin-by-barcode count matrix, removing barcodes with fewer than 1,000 unique fragments. We then cluster the barcodes using the graph-based Louvain algorithm using principal components as the input. Finally, we use *MACS2* to call peaks on the aggregated data for each cell cluster. The final set of peaks are generated by merging peaks less than 200 bp apart identified from different cell clusters.

#### Cell calling

As default, we use the filtering strategy to distinguish cell barcodes from non-cell barcodes, because the method is intuitive, easy to interpret and widely used among published studies [5]. We define a barcode as a cell if its total number of unique fragments is greater than 5,000 and the fraction of such fragments in peaks is greater than 50%. Users can use different thresholds for the fraction of fragments in enhancers, promoters or mitochondrial genome to filter barcodes as well.

#### Normalization

We provide two normalization methods. The Term Frequency-Inverse Document Frequency (TF-IDF) method [2,5] treats count data as binary and normalizes data by sequencing depth per cell and total number of unique fragments per peak. In the second method, count data is first log-transformed, followed by a linear regression to remove the confounding factor due to varying sequencing depth per cell for every peak. This method enables users to work with non-binarized count data. The TF-IDF method is set as the default normalization method in scATAC-pro.

#### Dimension reduction and data visualization

We use principal component analysis (PCA) as the default dimension reduction method because it is the most widely used method for scCAS data and easy to interpret. Note that if the PCA is conducted on the TF-IDF normalized data, such dimension reduction is also referred as *Latent Semantic Indexing* or *LSI* [2]. We use the PCA implementation in *Seurat* v3 with some modifications. In *Seurat* v3, the raw count matrix is log-transformed followed by a regression to remove confounding factors (the total number of unique fragments per cell). PCA is then performed on the transformed features. Because there are usually hundreds of thousand peaks in a scCAS data set, this process takes about a couple of hours to finish for a typical scATAC-seq data set. In our implementation, we first perform PCA on the normalized peak-by-cell count matrix, followed by a regression analysis on each principal component. This procedure substantially reduces the computation time and produces very similar clustering result as the original *Seurat* implementation (Supplementary Figure 1). Uniform Manifold Approximation and Projection (UMAP) [28] is used as the default visualization method.

#### Clustering

We provide the graph-based Louvain algorithm implemented in *Seurat* v3 as the default clustering method. Shared neighbor network (SNN) graph is constructed based on the first 30 principle components. Louvain algorithm is then performed on the SNN graph with default setting. We found that Louvain algorithm has a better balance between accuracy and running speed among several popular clustering methods (Supplementary Figure 2).

### scATAC-pro provides summary reports and interface to visualization tool

scATAC-pro generates global and cell-level quality assessment metrics for both aggregated and single cell data. Two types of metrics are generated. The first type of metrics evaluates data quality internally, including read mapping rate, duplicate rate, high-confidence mapped fragments (MAPQ greater than 30), and library complexity. The second type of metrics evaluates data quality using external annotations of genomic features, including fraction of fragments in mitochondrial genome, fraction of fragments overlapping with peaks and other annotated genomic regions, such as enhancers and promoters. The quality assessment summary reports are generated in html format. These statistics can be used to filter low quality barcodes.

Besides quality assessment metrics, scATAC-pro also generates summary reports for downstream analyses, including dimension reduction, clustering, differential accessibility analysis, TF motif enrichment analysis and footprinting analysis, gene ontology analysis, linking regulatory DNA sequences with gene promoters, and chromatin interactions prediction.

To enable interactive exploration of data in scATAC-pro, we provide an interface to *VisCello* [19], a visualization tool for single-cell omics data. To do this, we annotate the peaks with its nearest gene, and mark genes with their TSSs located within the peak. Users can then visualize chromatin accessibility signal of each peak or gene, and identify differential accessible peaks across different cell clusters.

### scATAC-pro provides utility functions to facilitate downstream analyses

#### Generation of input files for genome browser tracks for each cluster

It is a common task to visualize scCAS signal for each cell population on a genome browser. To generate normalized signal track file, bam file of cells from each cluster is first split from the bam file of all barcodes. Reads per cluster are then normalized as reads per kilobase per million mapped reads. scATAC-pro outputs normalized chromatin accessibility for each cluster in bigWig and bedGraph file formats, which can be directly uploaded to a genome browser for visualization.

#### Transcription factor footprinting analysis

ATAC-seq and related technologies use the Tn5 enzyme to cleavage nucleosome-free DNA while keeping the transcription factor binding sites intact due to protection by the bound TF. As a result, a small region, referred to as the footprint, exhibits reduced Tn5 cleavage rate at the ATAC-Seq peak locus. Unlike DNA motif analysis, TF footprinting analysis provides direct evidence of TF binding to the chromatin [29]. With Hint-ATAC [30], scATAC-pro enables footprinting analysis of either one cell cluster or differential TF binding between two cell clusters.

#### Integration of multiple scCAS datasets

Starting with bam files of multiple datasets, scATAC-pro first call peaks for each dataset. Peaks that are less than 200 bp apart are merged. Using this merged set of peaks based on all datasets, scATAC-pro generates raw peak-by-barcode count matrix, performs quality assessment, cell calling and downstream analysis for each dataset. Because we generate the peak-by-cell count using the same set of peaks for all samples, it is straightforward to integrate the datasets using existing tools, such as *Seurat* v3 or *Liger* [31]. scATAC-pro uses *Seurat* v3 as the default for this task.

#### Peak annotation and Gene Ontology analysis

To facilitate Gene Ontology analysis of genes associated with differential accessibility peaks, scATAC-pro first annotates each peak with its nearest gene. Gene Ontology analysis for those genes of each cluster can then be performed using the *runGO* module. This analysis helps users to further explore the identity of each cell cluster.

#### Predicting chromatin interactions by Cicero

Connecting regulatory DNA elements to target genes is a prerequisite to understanding transcriptional regulation. *Cicero* [32] predicts the interactions between cis-regulatory elements and the target genes using scCAS data. scATAC-pro generates the predicted interactions by running the module *runCicero*. The resulting interactions can be viewed through the UCSC genome browser.

## Case study

We used scATAC-seq data from 10,000 peripheral blood mononuclear cells (PBMCs) from a healthy donor (https://support.10xgenomics.com/single-cell-atac/datasets) to demonstrate the utility of scATAC-pro. Starting from the fastq files, scATAC-pro first demultiplexed sequencing reads by adding the cell barcodes (R2.fastq.gz) information to the paired-end reads (R1.fastq.gz, R3.fastq.gz). Adaptor sequences were then trimmed off, mapped to the GRCh38 reference genome using scATAC-pro default settings. Fragments with mapping quality score (MAPQ) less than 30 were removed. A summary report for mapping statistics and library complexity is provided for all reads (Figure 2A) and reads belonging to called cells (Figure 2B,C).

Using the default peak caller, scATAC-pro called 129,049 peaks after removing peaks overlapping with ENCODE blacklisted genomic regions (http://mitra.stanford.edu/kundaje/akundaje/release/blacklists/hg38-human/hg38.blacklist.bed.gz). Cell barcodes were selected by filtering out barcodes with fewer than 5,000 total unique fragments and the fraction of unique fragment in peak less than 50% (Figure 3A). Quality assessment report for each barcode was generated using various metrics, including distribution of insert size, transcriptional start site (TSS) enrichment profile, distribution of the total number of unique fragments for cell and non-cell barcodes, fractions of unique fragments overlapping with annotated genomic regions (Figure 3B-E). Overall statistics of data aggregated from all called cells was also computed (Figure 3F).

**Figure 3.**
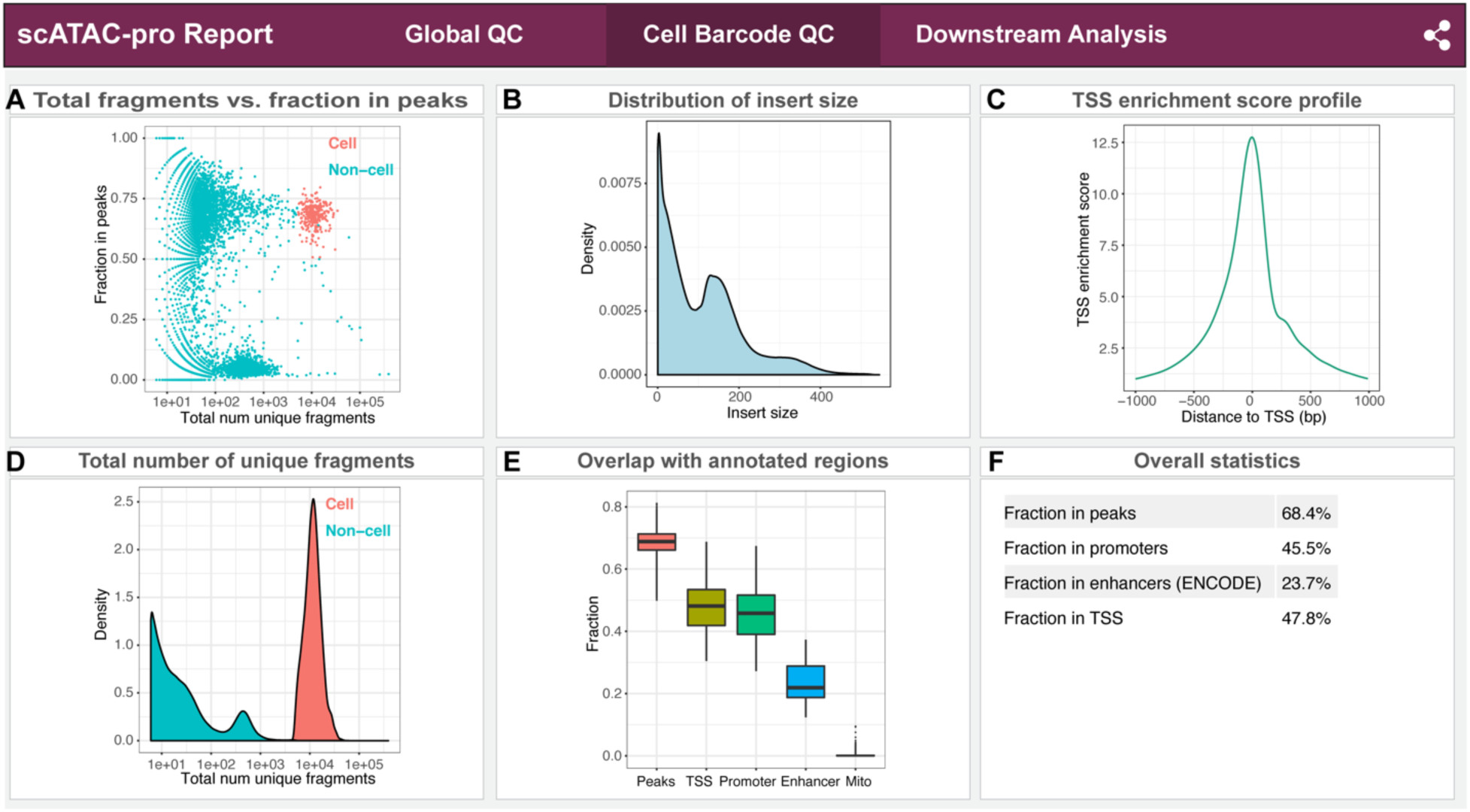
Quality assessment metrics for called single cells. scATAC-Seq data of human PBMCs was used for illustration purpose. (**A**) Plot of the fraction of fragments in peaks versus the total number of unique fragments. The plot can be used to distinguish cell barcodes from non-cell barcodes. (**B**) Distribution of insert fragment sizes. The plot can be used to evaluate the quality of transposase reaction. (**C**) Transcription start site (TSS) enrichment profile. (**D**) Distribution of the total number of unique fragments for cell and non-cell barcodes. The plot can be used to evaluate the amount of cell debris sequenced. (**E**) Boxplot of fragments overlapping annotated genomic regions per cell. (**F**) Overall statistics of data aggregated from all called cells.

Downstream analyses including clustering, TF motif enrichment analysis, TF footprinting analysis, and GO analysis, cis-element interaction prediction were conducted using default scATAC-pro methods and settings (Figure 4). In total, we found 9 cell types. The top 10 enriched TFs for each cluster are shown in Figure 4B, which provides a means for identifying cell type associated with each cluster. For example, binding motifs of PU.1(encoded by *SPI1*), IRF4, CEBPA, and CEBPB are highly enriched in clusters 0, 6, 7 and 8, suggesting those clusters are probably monocytes [33]. Motifs of EOMES and TBX5 were enriched in clusters 1, 2 and 5, suggesting those clusters are T cells. Enrichment of EBF1 [34] and BCL11A motifs [35] suggests cluster 3 represents B cells. The differential footprinting analysis between cluster 0 and cluster 1 further suggests that cluster 0 represents monocytes, because the monocytic TFs PU.1, JUNB, JUN, CEBPA and CEBPE [33,36,37] all have significant higher binding probability in cluster 0 cells (Figure 4).

**Figure 4.**
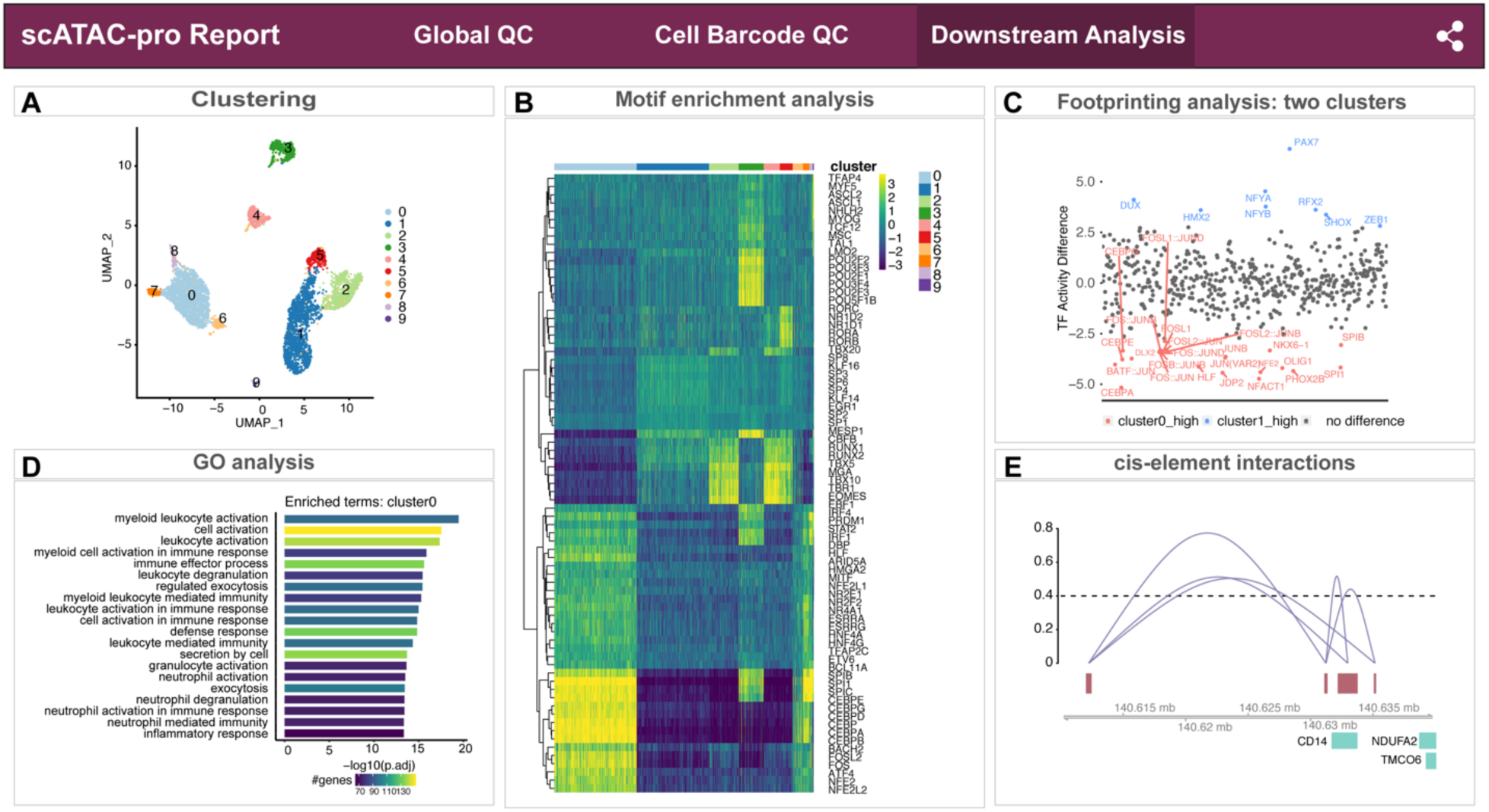
Summary report for downstream analyses. scATAC-Seq data of human PBMCs was used for illustration purpose. Results of the following analyses are shown: clustering analysis (**A**), transcription factor (TF) motif enrichment analysis (**B**), differential footprinting analysis between cluster0 and cluster1 (**C**), enriched gene ontology (GO) terms for cluster0 (**D**), and predicted cis-interactions at *CD14* locus (**E**).

Using *VisCello [19],* we can display chromatin accessibility values of TSS regions of several marker genes across cell clusters (Figure 5A and Supplementary Figure 4A), such as *MS4A1* (CD20) for B cells, *GNLY* and *NKG7* for natural killer (NK) cells, *CD3E* for T cells, *CD14, LYZ*, and *FCGR3A* (CD16) for monocyte cells, and *CST3* for dendritic cells (DC) [38]. We also displayed UCSC genome browser tracks for two example genes, *CD14* (Figure 5B) and the *FCER1A* (Supplementary Figure 4B). Taken together, based on the chromatin accessibility profile of known cell-type-specific marker genes, we annotated cell cluster as T cells, B cells, CD14^+^ monocytes, CD16^+^ monocytes, dendritic cells, natural killer cells.

**Figure 5.**
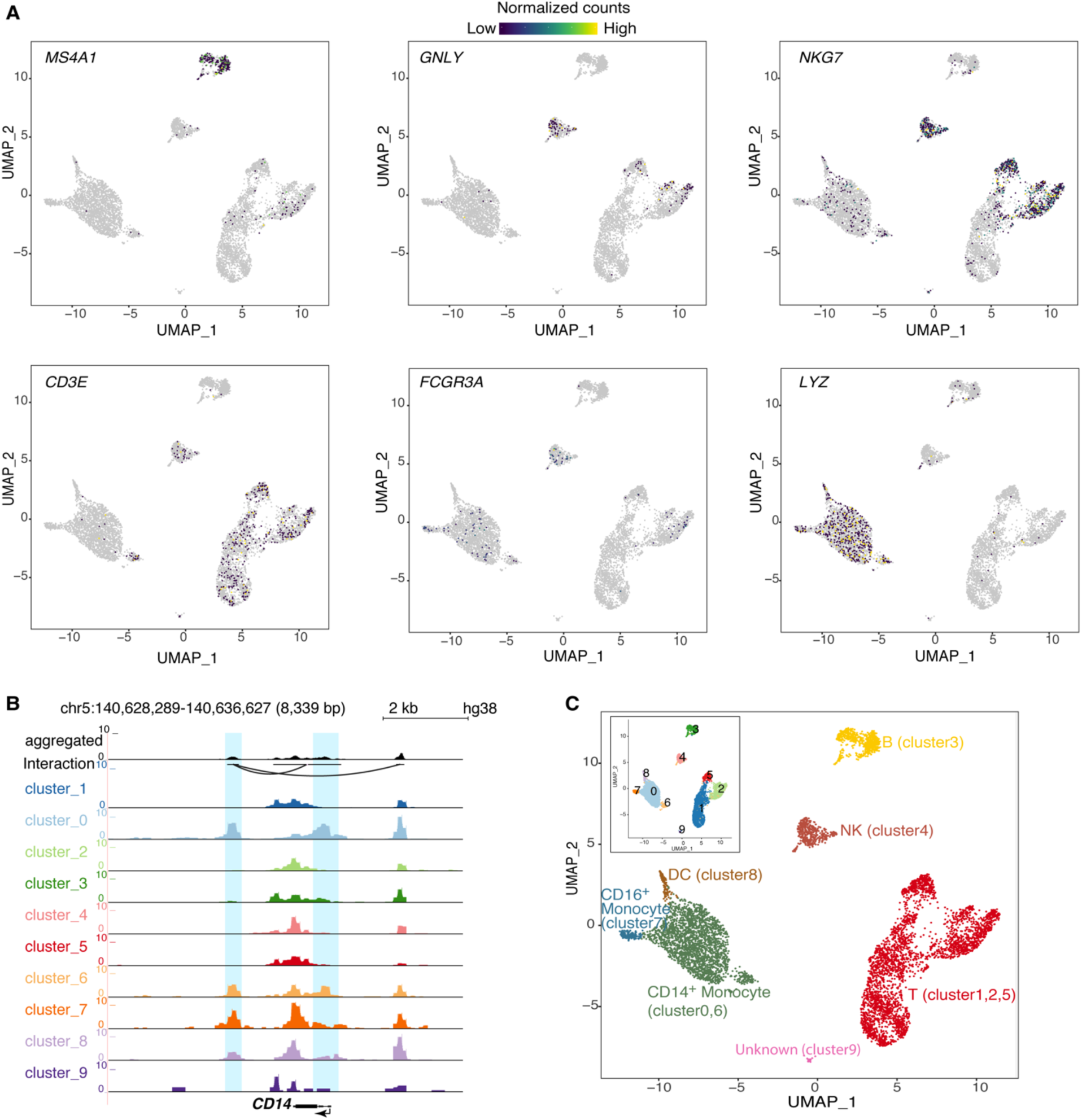
Visualization of scATAC-Seq data. (**A**) Chromatin accessibility signal of single cells. Normalized chromatin accessibility signal for peaks overlapping with transcriptional start sites of selected marker genes. Data is visualized using visCello. (**B)** Chromatin accessibility signal of aggregated cells along the genome. Genome browser view of normalized chromatin accessibility signal at the CD14 locus across cell clusters. (**C)** Cell type assignment based on chromatin accessibility signals of known cell type marker genes. Inset, clustering result without cell type assignment.

## Discussion

scATAC-pro provides a comprehensive solution for scCAS data QC and analysis. It reports a number of commonly used QC metrics for both aggregated data of all barcodes and barcodes of called cells. These metrics evaluate multiple steps of the experimental protocol, including the transposase reaction (insert size), quality of nucleus preparation (insert size, fraction of unique reads, cell vs non-cell reads, mitochondrial reads), cell encapsulation (cell vs non-cell reads), PCR amplification (duplicate rate), library preparation and sequencing (fraction of unique reads and reads with MAP > 30). Although there is no universally optimal QC metric for all kinds of scCAS data, the fraction of fragments in peak per cell is the most widely used in the literature. Alternative metric such as the TSS enrichment score per cell is introduced recently [17], but its utility may be limited for cell types that have a large fraction of active TSS-distal peaks. Having a comprehensive annotation of cis-regulatory elements across all human cell types will facilitate the task of evaluating quality of gene-distal ATAC-Seq peaks.

Because there is no clear optimal method for many analytical tasks, scATAC-pro provides multiple methods that allow users to tailor their analyses and to address dataset-specific characteristics. To guide the users, we have provided carefully evaluated default setting for each analytical task, including both method choice and parameter setting of the selected method(s).

The open-source and modular design of scATAC-pro facilitates the maintenance and future development of the software. Several experimental protocols exist for generating scCAS data. Data generated using these protocols have different characteristics and qualities. scATAC-pro is the first software tool that enables analysis of all types of scCAS data. By doing so, scATAC-pro facilitates the integration of rapidly growing scCAS data.

The current version of scATAC-pro generates a static summary report. It can be enhanced by generating dynamic summary report in future versions of the software. For example, for downstream analyses, the results can be updated in real time based on the cell clusters compared. Display of cis-interactions within an arbitrary genomic region specified by the user will be another very useful feature to add.

## Conclusions

scATAC-pro is a comprehensive open source software tool for processing, analyzing and visualizing single-cell chromatin accessibility sequence data. With the rapid accumulation of single-cell chromatin accessibility sequencing data, application of scATAC-pro will facilitate a better understanding of epigenomic heterogeneity in healthy and diseased tissues.

## Methods

### scATAC-pro workflow

scATAC-pro consists of two units, data processing unit and downstream analysis unit (Figure 1). The data processing unit takes raw fastq files for reads and barcodes as the input and outputs peak-by-cell count matrix, QC report and genome track files. It consists of the following modules: demultiplexing, adaptor trimming, read mapping, peak calling, cell calling, genome track file generation and quality control assessment. The downstream analysis unit consists of the following modules: dimension reduction, cell clustering, differential accessibility analysis, Gene Ontology analysis, TF motif enrichment analysis, TF footprinting analysis, linking regulatory DNA sequences with gene promoters, and integration of multiple datasets. We provide flexible options for all modules.

We designed the workbench to be user friendly. In each run, users just need to specify the input file (“*--input*” flag), the module name (“*--step*” flag) and a configuration file *(“--config”* flag), in which users provide additional parameters or options for the analysis modules. Users can choose to run the entire or partial workflow. By default, all results are saved in the “output” directory under the current directory *(--output_dir “./output”*).

#### Demultiplexing

Given the fastq files and the barcode fastq files, the barcode sequences are written into the name of each read sequence (in the format as @BARCODE:ORIGINAL_READ_NAME) to facilitate the tasks of downstream modules, such as generating peak-by-cell matrix and quality assessment at single cells. For data generated using 10x genomics, sciATAC-seq and dsciATAC-seq protocols, users need to provide the paired-end read fastq files and the index fastq file (also supports multiple index fastq files). For scTHS-seq data, users need to specify the parameter *isSingleEnd=TRUE* in the configuration file because scTHS-seq data are single-end reads. This module is skipped if the barcode for each read is recorded in the required format. For example, in the mouse sci-ATAC-seq atlas data, [2,5], the barcode for each read is saved in the name of each read in the SRA file.

#### Adaptor trimming

To map sequencing reads confidently to the reference genome, scATAC-pro first trims off the adaptor sequence and primer oligo sequence from raw reads using *trim_galore (http://www.bioinformatics.babraham.ac.uk/projects/trim_galore/)* as the default, which can automatically detect and trim the adaptor and primer sequences. Alternatively, users can also use *Trimmomatic* [39], which is faster but users need to specify the sequence of the adaptor in the configuration file.

#### Read mapping

Different alignment methods make different compromise between mapping accuracy and speed. BWA, Bowtie and Bowtie2 are three popular and top-ranked aligners based on previous benchmarking studies [20,27]. scATAC-pro enables all three aligners for read mapping. Based on its popularity for single cell ATAC-seq studies, cellranger-atac use BWA as its aligner, we use BWA (bwa-mem) as the default aligner based on its balance between mapping speed and accuracy. Users can provide addition options in the configuration file by specifying *MAPPTING_METHOD* and corresponding parameters. For instance, if users want to use 10 CPUs for parallel computing, they can set *BWA_OPTS = -t 10* if BWA is used, *BOWTIE_OPTS = -p 10* if BOWTIE is used and *BOWTIE2_OPTS = -p 10* if Bowtie2 is used. After mapping, scATAC-pro uses samtools [40] to sort, index, mark duplicates and filter low quality reads in the bam file.

The position sorted bam file, filtered bam file (default MAPQ score > 30), and the mapping statistics are automatically generated and saved for downstream modules. A file called *fragment.txt* that records the genomic location, barcode and the number of duplicates of each unique fragment is generated using a custom R script to facilitate downstream analysis.

#### Peak calling

By default, we identify open chromatin regions by identifying peaks using aggregated fragments across all barcodes. *MACS2* is a popular peak calling tool for ATAC-Seq and ChIP-Seq data. We also enable the *GEM* algorithm for peak calling. It is recommended by the ENCODE consortium for its good performance on calling TF motif enriched peaks. The processed scCAS data is then summarized as the peak-by-barcode matrix. Peaks that only appear in rare cell types are challenging to call by the above approach of using aggregated fragments across all barcodes. An alternative approach is to bin the genome without peak calling or combination of binning the genome and peak calling [5]. For the combination strategy, we first segment the genome into 5 kb bins and generate bin-by-barcode count matrix, removing barcodes with low total number of fragments (e.g. 1,000). We then cluster the barcodes followed by peak calling for each cluster. Peaks or bins overlapped with blacklisted genomic regions are removed for downstream analysis. Users can specify *PEAK_CALLER* to be one of *MACS2, GEM, BIN, or, COMBINED* in the configuration file.

#### Cell calling

Not all barcodes are real cells in a typical scCAS dataset due to cell collision and/or cell debris. How to distinguish cell barcodes from non-cell barcodes is still a challenging problem. Generally, users select cell barcodes either by filtering out low quality barcodes based on some summary statistics, such as total number of fragments and fraction of fragments in peak regions. Alternatively, users can use model-based approaches. For example, the *cellRanger-atac* method fits a mixture of two zero-inflated negative binomial models to discriminate cell barcodes and non-cell barcodes. *EmptyDrops* [41], originally designed to identify cells from scRNA-Seq data, models the counts using a Dirichlet-multinomial distribution. scATAC-pro provides all of the aforementioned strategies/methods. Based on our experience, cellranger-atac and *EmptyDrops* (with the default fdr of 0.001) tend to call too many cells, while the knee point approach of *EmptyDrops* and *cellRanger-atac* are too stringent. Therefore, we choose the filtering strategy as the default since it is simple and intuitive. For the filtering strategy, users can filter barcodes based on one or multiple summary statistics such as the total number of unique fragments, fraction of fragments in peaks, fraction of fragments in mitochondrial genome, and fraction of fragments overlapping with annotated promoters, enhancers, and TSS regions. Since the implementation of *cellRanger-atac* cell calling is not publicly available, we implemented the algorithm using custom R scripts.

#### Quality assessment

scATAC-pro provides mapping statistics for all reads as well as reads belonging to called cells. The following QC metrics are reported: total reads, total number of mapped reads, unique mapping rate, fraction of reads in mitochondrial genome, number of duplicate reads, high-quality reads (MAPQ >30), library complexity, fraction of reads in annotated genomic regions and TSS enrichment profile. The same set of summary statistics is also reported for reads belonging to called cells. scATAC-pro also reports the number of cells called, median number of fragments per cell, fraction of mapped reads belonging to cells.

#### Normalization

The default Term Frequency-Inverse Document Frequency (TF-IDF) normalization is implemented using the TF.IDF function in *Seurat* v3. We also provides an alternative normalization method, which first log-transforms the count followed by regression to remove the confounding factor due to sequencing depth per cell.

#### Dimension reduction, cell clustering, and visualization

scATAC-pro supports principal component analysis (PCA) (which is also called Latent Semantic Indexing (LSI) if the data were first normalized by TF-IDF) and latent Dirichlet allocation (LDA) for dimension reduction. We use the *Seurat* v3 toolkit to implement PCA, Louvain clustering algorithm, and the R CisTopic package to implement LDA. We provide t-distributed stochastic neighbor embedding (tSNE) and uniform manifold approximation and projection (UMAP) (implemented by *Seurat* v3) for visualization.

#### Differential accessibility analysis

Peaks with differential accessibility across different cell clusters are potentially cell-specific gene regulatory elements. We use Wilcoxon test as the default method to perform differential accessibility analysis. Alternative methods such as logistic regression based (LR) method (implemented in *Seurat* v3), DESeq2 [42] and negative binomial regression based test (implemented in *Seurat* v3) are also available. Users can compare any given two clusters or compare one cluster versus the rest of the clusters using the module *runDA* and specify *group1, group2* in the configuration file.

#### Generation of genome browser track files

scATAC-pro outputs bigWig and bedGraph files for visualizing chromatin accessibility signal in a genome browser. The signal is normalized by reads per kilobase per million mapped reads, for aggregated data or for each cell cluster. Those files are generated using the *bamCoverage* command in *deepTools* toolkit [43].

#### TF motif enrichment analysis

scATAC-pro constructs the TF binding accessibility profile for each single cell using the *chromVAR* with a slight modification. *chromVAR* computes a gain or loss accessibility score for peaks sharing the same motif by comparing to accessibility score of peaks with similar mean accessibility and GC contents. To speed up this analysis, in scATAC-pro, instead of using the whole peak-by-cell matrix, we select top 30% of most variable peaks. This reduces the running time of *chromVAR* by eight times compared to using the full matrix of the PBMC data. We then identify TFs that have significantly higher accessibility in one cell cluster than in the other cell cluster by conducting a two-sample Wilcoxon test. TFs that are significantly higher chromatin accessible in each cell cluster is saved in a text file and visualized using heatmap.

#### TF footprinting analysis

We use Hint-ATAC [30] to perform TF footprinting analysis, which is the first tool designed specifically for ATAC-seq data. Due to the sparsity of scCAS data, it is impossible to predict TF footprints at single cell level, but feasible at cell cluster level since the read depth per cluster is similar to bulk ATAC-seq data. We also use Hint-ATAC to do differential TF footprinting analysis.

#### Summary reports

scATAC-pro automatically updates the summary report after processing and/or downstream analysis were done through custom R scripts. If some analysis modules are not executed, scATAC-pro still generates the report with results of executed modules.

### Benchmarking of clustering algorithms for scCAS data

The single-cell data were simulated by resampling bulk ATAC-seq data on 13 primary human blood cell types [7]. Specifically, we simulated data for 200 cells for each of the 13 cell types. For each cell, 10,000 reads were randomly selected from the mapped reads in the bulk data. Peaks were called using MACS2 using the aggregated single-cell data. The performance of each method was evaluated using the adjusted rand index and bulk sorted cell types as the ground truth (Supplementary Fig 2). To investigate the robustness of each method as a function of cell type composition, we sampled a total of 1,000 cells from the 13 cell types with different cell type compositions. The fractions of different cell types were generated based on the Dirichlet distribution (with shape parameter alpha = 3 for each component). For each method, the default parameters were used, except for the number of clusters, which was set to 13.

## Availability of data and materials

The datasets analyzed during this study are included in this published article and its supplementary information files. The software packages are available from the GitHub repository https://github.com/tanlabcode/scATAC-pro under a MIT license and are also deposited in zenodo with DOI: 10.5281/zenodo.3520939.

## Authors’ contributions

WY and KT conceived and designed the study. WY designed and implemented the scATAC-pro software with the help of YU, QZ, and CC. YU and QZ provided additional analytical tools. WY performed data analysis. KT supervised the overall study. WY and KT wrote the paper.

## Acknowledgements

We thank the Research Information Services at the Children’s Hospital of Philadelphia for providing computing support.

---

**Table 1.**
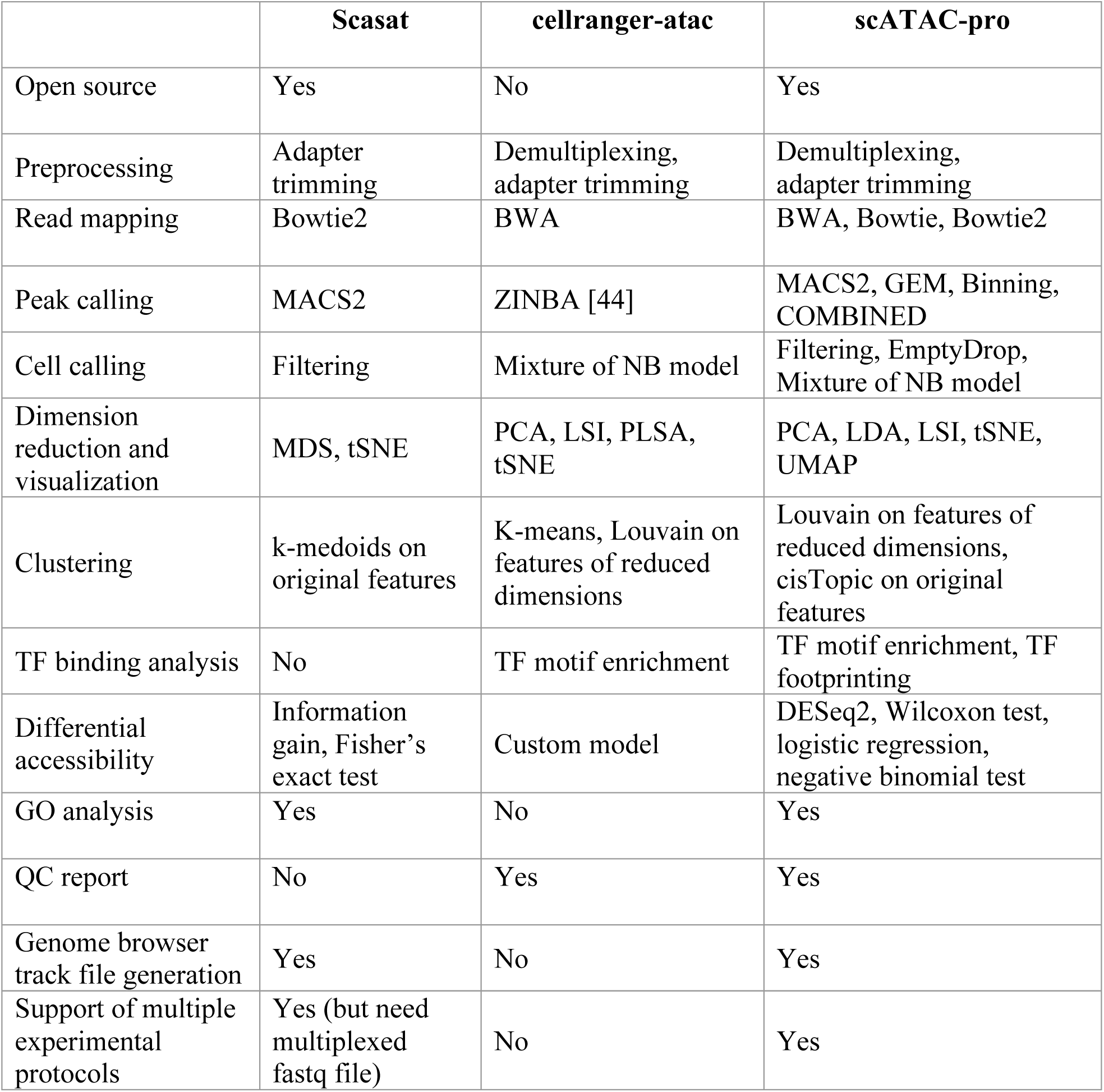
Comparison of the features of three comprehensive software for the processing and analysis of single-cell chromatin accessibility sequencing data. MDS: multidimensional scaling; NB: negative binomial; PLSA: probability latent semantic analysis; LSI, latent semantic indexing; LDA, latent Dirichlet allocation; UMAP, uniform manifold approximation and projection (UMAP); tSNE, t-distributed stochastic neighbor embedding.

|  | **Scasat** | **cellranger-atac** | **scATAC-pro** |
| --- | --- | --- | --- |
| Open source | Yes | No | Yes |
| Preprocessing | Adapter trimming | Demultiplexing, adapter trimming | Demultiplexing, adapter trimming |
| Read mapping | Bowtie2 | BWA | BWA, Bowtie, Bowtie2 |
| Peak calling | MACS2 | ZINBA [44] | MACS2, GEM, Binning, COMBINED |
| Cell calling | Filtering | Mixture of NB model | Filtering, EmptyDrop, Mixture of NB model |
| Dimension reduction and visualization | MDS, tSNE | PCA, LSI, PLSA, tSNE | PCA, LDA, LSI, tSNE, UMAP |
| Clustering | k-medoids on original features | K-means, Louvain on features of reduced dimensions | Louvain on features of reduced dimensions, cisTopic on original features |
| TF binding analysis | No | TF motif enrichment | TF motif enrichment, TF footprinting |
| Differential accessibility | Information gain, Fisher's exact test | Custom model | DESeq2, Wilcoxon test, logistic regression, negative binomial test |
| GO analysis | Yes | No | Yes |
| QC report | No | Yes | Yes |
| Genome browser track file generation | Yes | No | Yes |
| Support of multiple experimental protocols | Yes (but need multiplexed fastq file) | No | Yes |

## Supplementary figure legend

**Supplementary Figure 1.**
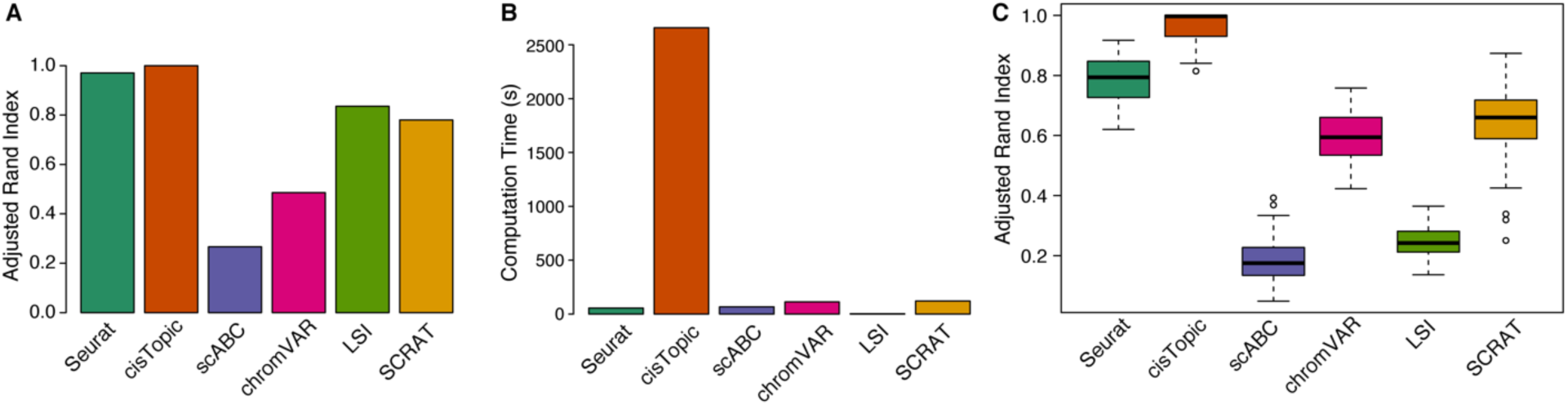
Performance comparison of principal component analysis (PCA) implemented in *Seurat* and scATAC-pro. **(A)** Computation time as a function of the fraction of features (peaks) used. **(B)** Similarity of the clustering results based on PCA by *Seurat* and scATAC-pro. Clustering was done using the Louvain algorithm. Similarity was measured using the adjusted rand index.

**Supplementary Figure 2.**
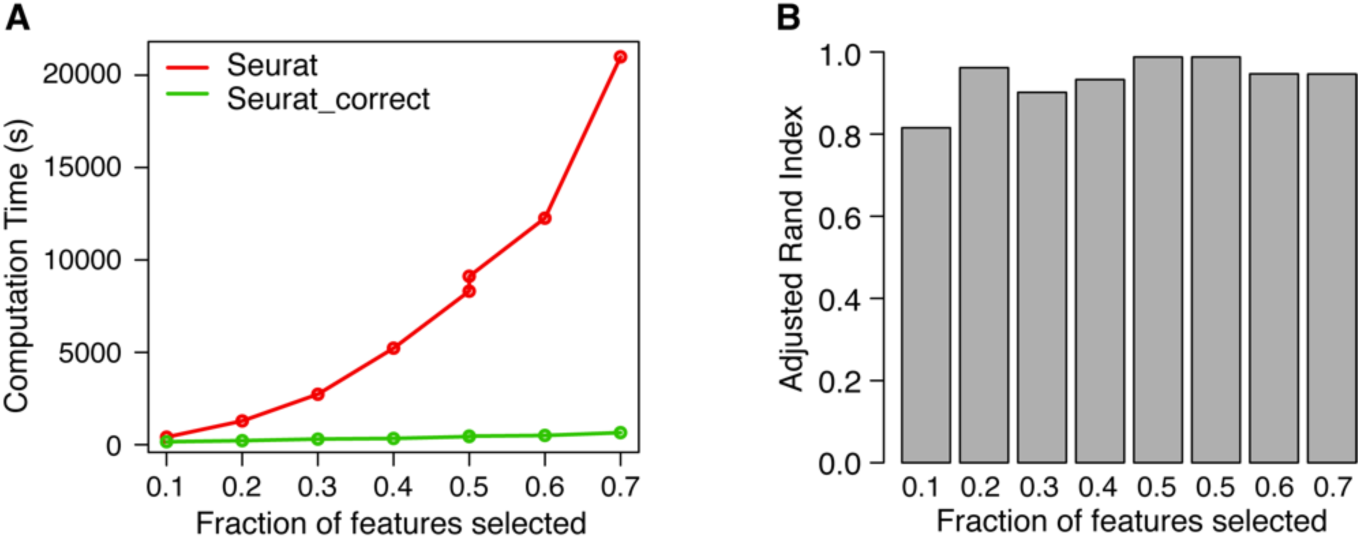
Performance comparison of different clustering algorithms using simulated data. **(A)** Adjusted rand index for different algorithms using the FACS-sorted cell types as the ground truth. Cells from 13 types were sampled with equal probability. **(B)** Computation time of each method. **(C)** Adjusted rand index of 100 sets of simulated data. Cells were sampled from the 13 types with different proportions. The proportions of different cell types were generated based on the Dirichlet distribution (with shape parameter alpha = 3 for each component).

**Supplementary Figure 3.**
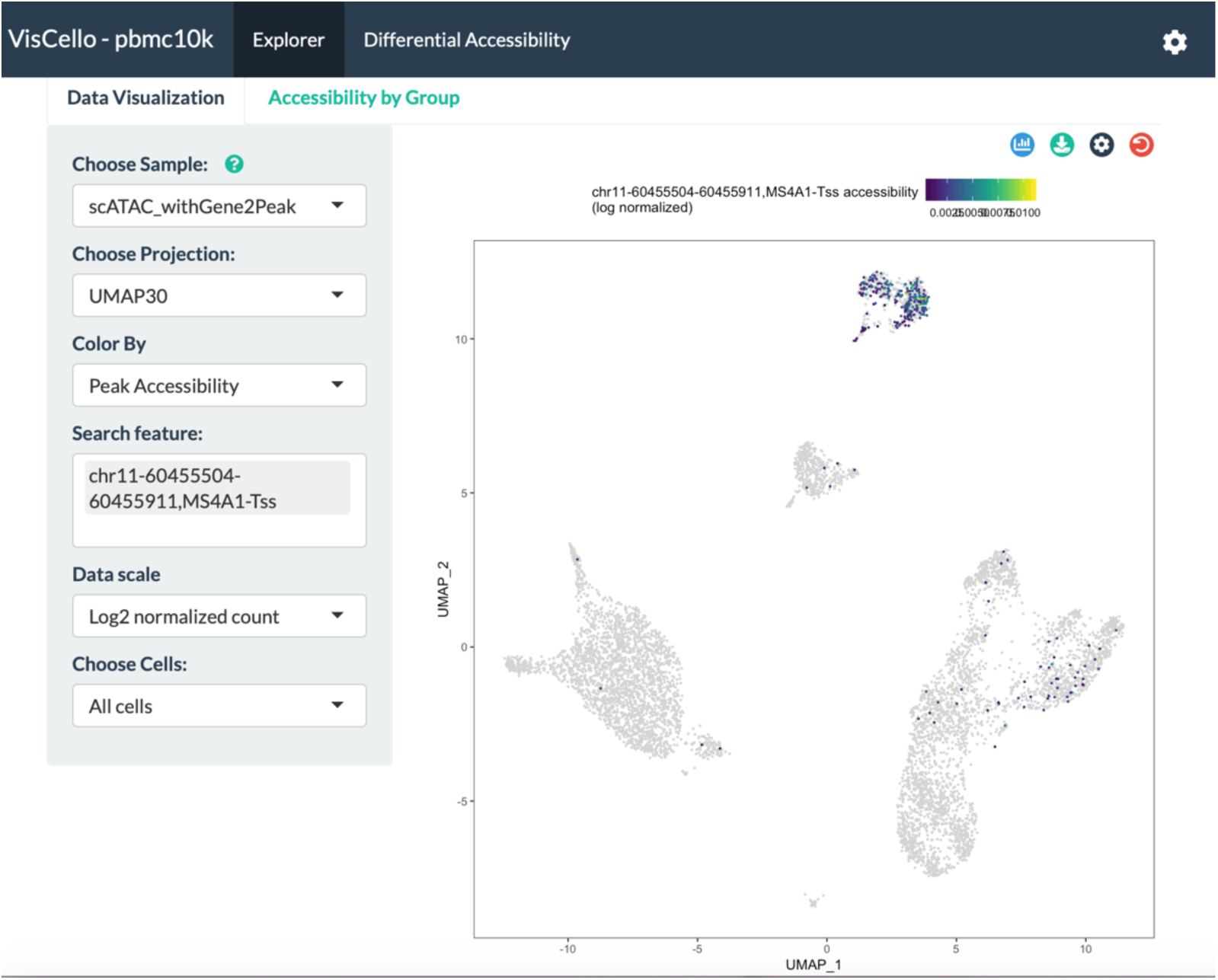
Screenshot of the user interface of the visualization tool, *VisCello.* scATAC-Seq data of human peripheral blood mononuclear cells (PBMCs) was used for illustration purpose. Chromatin accessibility score of peak overlapping with the transcriptional start site of *MS4A1* is displayed. Users can use gene name or peak coordinate as the search keyword to explore the accessibility of interested regions. The raw and normalized data can be visualized using uniform manifold approximation and projection (UMAP) or t-distributed stochastic neighbor embedding (tSNE) with different numbers of principal components

**Supplementary Figure 4.**
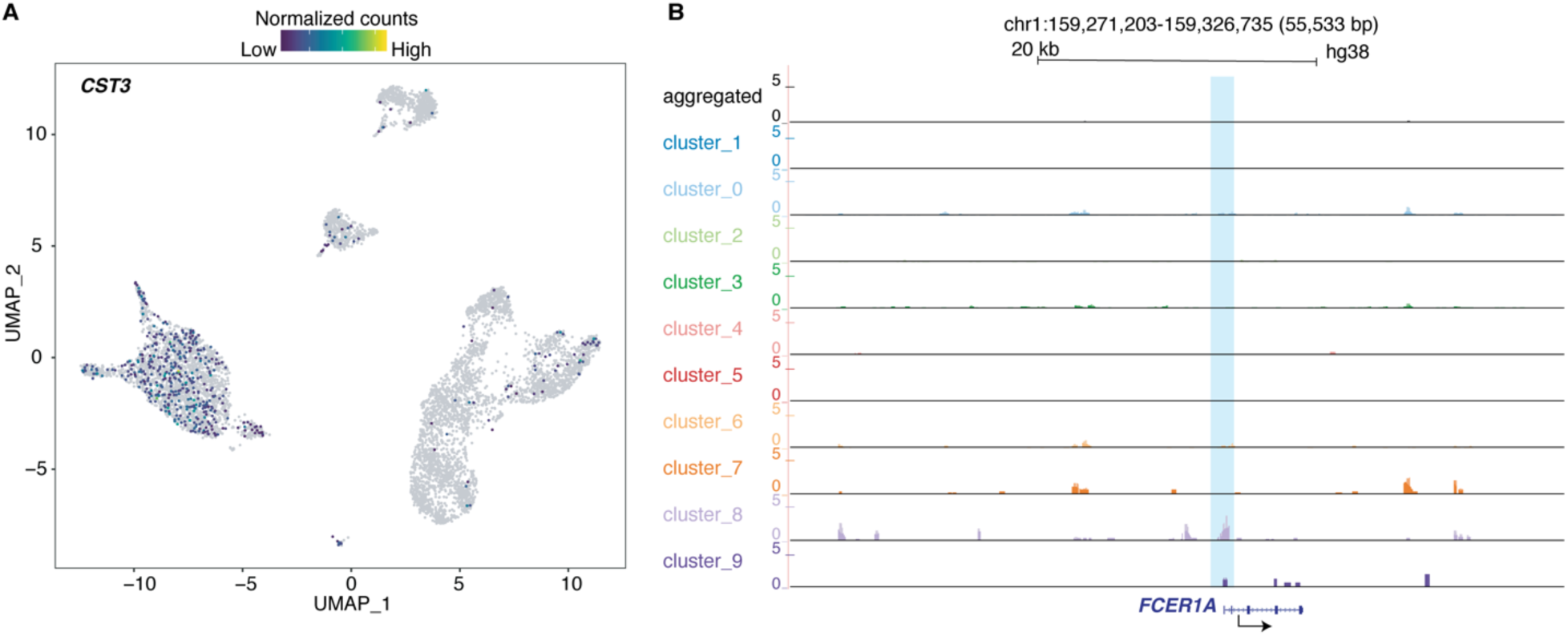
Chromatin accessibility of transcription start site (TSS) of two dendritic cell markers *CST3* (A) and *FCER1A* (B) shown in UMAP and UCSC genome browser, respectively.

